# Reproducing squat kinematics after total knee replacement with bi-level inverse optimal control

**DOI:** 10.64898/2026.06.11.731549

**Authors:** Hojin Song

## Abstract

Total knee replacement restores mobility in patients with advanced osteoarthritis, yet many individuals still experience limited ability to perform high-flexion tasks such as squatting. Current preoperative planning relies on static imaging and does not evaluate postoperative high-flexion kinematics. In this study, we developed a simulation framework that uses bi-level inverse optimal control to reproduce postoperative squat kinematics from subject-specific models.

Musculoskeletal models were built for six total knee replacement patients from experimental squat recordings. Cost-function weights were identified in an individualised setting and in a group-level setting. The former gives a subject-specific reconstruction when postoperative motion is available. The latter yields one shared cost function that does not need each patient’s postoperative trajectory. That shared function was then tested with leave-one-subject-out cross-validation.

The individualised setting reproduced experimental trajectories with low errors across all joints (mean apex difference 1.53^◦^, root-mean-square error 5.15^◦^, normalised root-mean-square error 11.15%, Pearson correlation 0.96). The in-sample group-level setting yielded higher errors (mean apex difference 5.70^◦^, root-mean-square error 6.75^◦^, normalised root-mean-square error 17.53%, Pearson correlation 0.95) while preserving the general pattern and phasing of the motion. Under cross-validation the shared cost function still worked on left-out subjects; the extra error was small when all joints are averaged (7.00^◦^, 7.36^◦^, 17.75%, and 0.95 for the same four metrics), although that loss was concentrated at the hip. The full stand-to-squat-to-stand cycle was generated in a single optimisation without prescribing a target squat depth.

A shared cost function can therefore still reproduce squat kinematics in patients who were left out of the fit. That is required before the method could be used preoperatively. It does not itself predict a squat from preoperative data. Larger and independent cohorts are needed before the method could be used to compare alignment choices.

## Introduction

Total knee replacement (TKR) is a well-established procedure for alleviating pain and restoring mobility in patients with advanced knee conditions such as osteoarthritis and rheumatoid arthritis [1]. While early TKR focused primarily on pain relief and correction of deformities, modern surgical techniques increasingly emphasise the restoration of functional capacity, allowing patients to resume daily activities such as squatting [2–4]. Despite these advances, 15–20% of TKR patients still report dissatisfaction with their outcome [5, 6].

Current preoperative planning relies largely on static imaging and templating to determine implant alignment. Surgeons commonly use radiographs or CT scans to position femoral and tibial components along mechanical or anatomical axes [7]. Although these approaches provide important information for static correction, they do not evaluate postoperative high-flexion kinematics. Squatting requires substantial knee flexion and is an important marker of recovery, yet planning workflows still centre on static alignment and component position [8, 9].

Squatting consists of distinct phases, including descent, deep flexion, and ascent, each of which places high demands on knee stability, quadriceps strength, and range of motion. Post-TKR patients may struggle with these phases due to altered joint mechanics, reduced quadriceps strength, and pain avoidance. Consistent with this, squatting has been reported as one of the most difficult functional activities for patients to perform after TKR [10]. Biomechanical indicators such as peak knee flexion angle are therefore important markers of functional recovery. However, there remains a need for computational frameworks that can reproduce high-flexion kinematics in TKR patients from a subject-specific model, as a prerequisite for later asking how surgical parameters affect that motion.

Optimisation-based motion generation has emerged as a promising approach to address this problem. These methods formulate human movement as the minimisation of a cost function under biomechanical constraints, allowing motion to be generated without requiring large training datasets [11, 12]. In contrast, data-driven approaches have shown success for activities such as squatting and running, but often depend on extensive patient-specific data that may be unavailable or difficult to collect in the TKR population [13–15]. Physics-based optimisation therefore provides a complementary route for generating squat kinematics from a subject-specific model when a large patient-specific training set is not available.

In earlier work we used a torque-driven optimal control model to generate squat trajectories while rotating the tibial component away from a neutral pose [16]. That study treated the cost-function weights as fixed and asked how alignment changed the predicted motion. It left open which weights should be used for TKR patients, and whether one set of weights can be reused. The present study addresses those questions. Subject-specific femoral and tibial orientations from the CAMS-Knee dataset [17, 18] are built into a 3-DOF knee model with ligament dynamics, and bi-level inverse optimal control is used to identify the weights from measured squat trajectories of six TKR patients, in an individualised setting, a group-level setting, and leave-one-subject-out cross-validation.

The aim of this study was to develop and preliminarily evaluate a bi-level inverse optimal control framework for reproducing post-TKR squat kinematics. The individualised setting measured how well subject-specific weights can match a recorded squat when postoperative motion is available. The group-level setting asked whether one shared cost function can still recover the main features of the squat across this cohort. Leave-one-subject-out cross-validation then re-estimated those group-level weights without each subject in turn and scored the held-out trajectory. The comparison is between fitting a cost function to a patient and applying a cost function that patient did not help identify.

## Materials and methods

Our workflow comprises three key steps. First, we developed subject-specific skeletal models using Open-Sim [19], parameterised to represent TKR component alignment. Each model was scaled to match the corresponding subject and configured using the available component alignment information. Next, using this parameterised model, squat motion was simulated by solving an optimal control problem (OCP) under the assumption that human movement can be represented as the optimisation of a weighted cost function, as shown in previous studies [11, 12, 20–22]. The squat was selected because it requires substantial knee flexion and is among the activities that TKR patients most frequently report difficulty performing after surgery [10].

Because the precise cost function governing human motor control cannot be directly measured, we employed a bi-level optimisation framework to identify the cost-function weights from experimental motion data. Three settings were investigated. The individualised setting identified subject-specific weights and was used as a reconstruction benchmark for the best achievable agreement with each subject’s observed motion. The in-sample group-level setting identified a single set of weights across all subjects, fitted and evaluated within the same cohort, and was used to examine whether a population-level cost function could reproduce the general squat kinematic patterns observed within that cohort. To evaluate whether this population-level cost function generalises to subjects not used in its derivation, we additionally performed leave-one-subject-out cross-validation, in which group-level weights were re-estimated excluding each subject in turn and applied to that subject’s own OCP. Finally, the simulated trajectories from all three settings were compared with experimental squat data from TKR patients obtained from the CAMS-Knee database [17, 18].

### Skeletal model

We employed a biomechanical model with 12 DOFs, adapted from the advanced knee flexion model developed by Catelli et al. [23], illustrated in Fig 1. This model encompasses 3 DOF in the pelvis (1 rotational and 2 translational), 3 DOF at the hip (all rotational), 3 DOF for knee joint movement (all rotational), 2 DOF for ankle motion (both rotational), and 1 DOF for lumbar extension (rotational).

**Fig 1.**
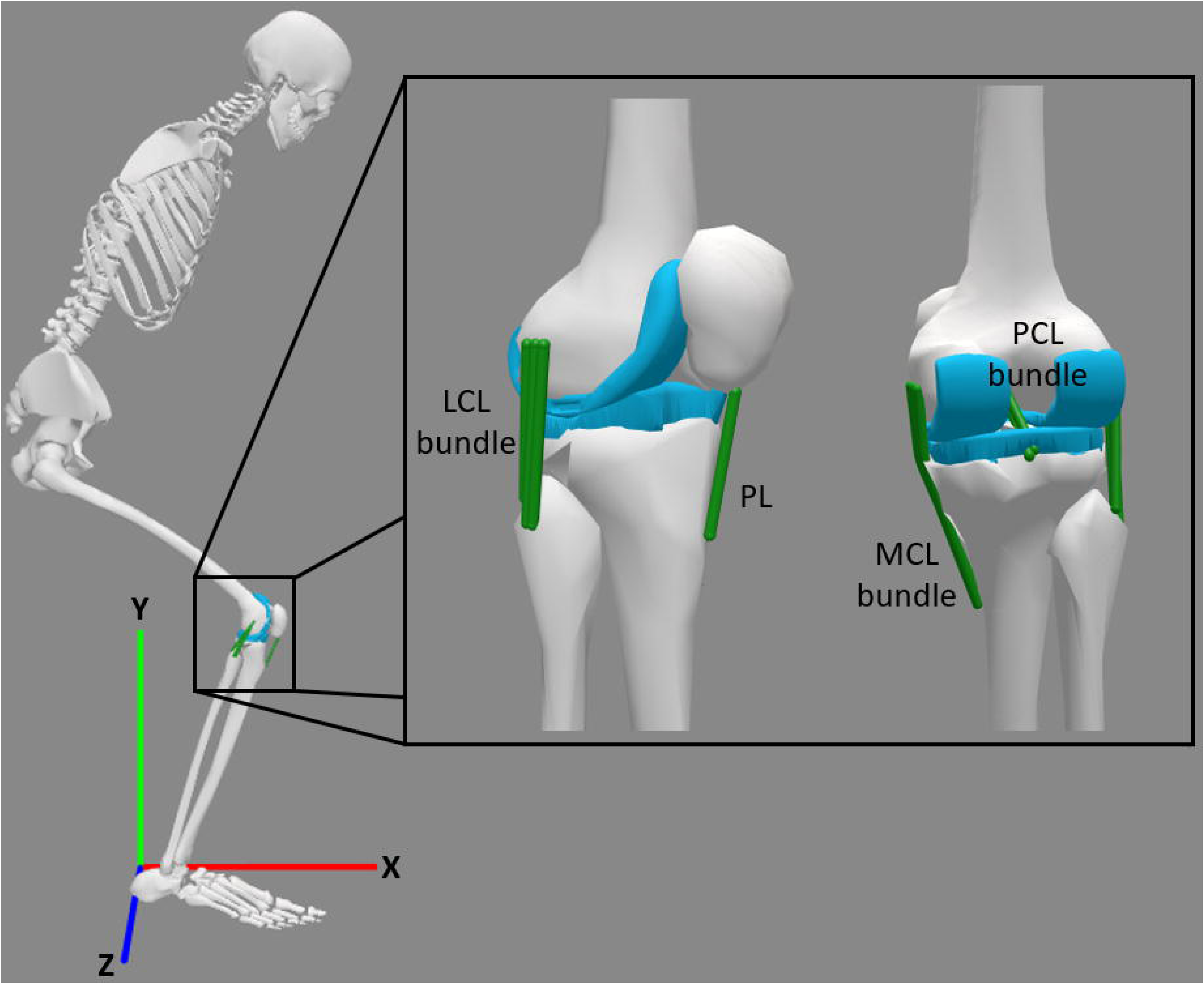
MSK model utilised in the study (left), along with the depicted positions of knee ligament components (right). The model encompasses distinct bundles for the medial and lateral collateral ligaments (MCL and LCL), as well as a bundle for the patellar ligament (PL) and posterior cruciate ligament (PCL). The patellar ligament (PL) is considered rigid within this representation.

The connection of the foot to the ground was modelled to maintain continuous foot-ground contact throughout the squat. Seven torque actuators were used to actuate the hip, knee, ankle, and lumbar joints. Soft tissue around the knee was represented by eight force-producing ligament bundles plus a rigid patellar ligament.

The ligaments were modelled as non-linear elastic forces to mimic the mechanical behaviour of anatomical knee ligaments, as proposed by Blankevoort [24].

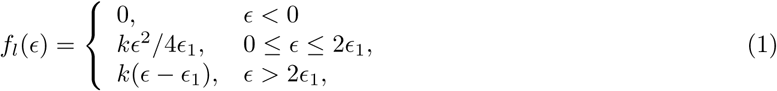

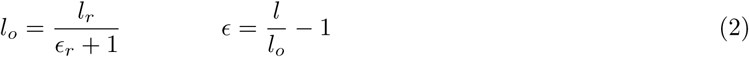

where *f_l_*(*ɛ*) is the force exerted by a ligament, *k* is its stiffness, and *ɛ* is the ligament strain. Here, *ɛ*_1_ was fixed at 0.03 [25] and defines the transition to the linear portion of the force-strain curve. The parameter *l* is the current ligament length, *l_r_* is its length at full knee extension, *l_o_* is the resting length, and *ɛ_r_* is the reference strain at a specified reference position. The stiffness parameters and reference strain values for each ligament are provided in Table 1 [24, 26].

**Table 1.** Stiffness and reference strain of each ligament [24, 26].

| **Ligament** | **Stiffness, k (N)** | **Reference Strain,** ϵ |
|---|---|---|
| aPCL | 9000 | -0.24 |
| pPCL | 9000 | -0.03 |
| aLCL | 1667 | 0.03 |
| mLCL | 1667 | 0.03 |
| pLCL | 1667 | 0.03 |
| aMCL | 2750 | 0.04 |
| mMCL | 2750 | 0.04 |
| pMCL | 2750 | 0.05 |

Subject-specific TKR component orientations were assigned in the MSK model. Fig 2 illustrates key geometric aspects of the knee joint and the positioning of the knee replacement. In Fig 2(a), the Hip-Knee-Ankle (HKA) angle is shown, defined as the angle between the mechanical axes of the femur and tibia in the coronal plane. Fig 2(b) presents the posterior tibial slope (PTS) in the sagittal plane, defined as the slope of the tibial plateau relative to the sagittal mechanical axis of the tibia. This angle is important for anterior-posterior knee stability [27]. Fig 2(c) and (d) depict the varus-valgus alignments of the femoral and tibial components, respectively. The angles in Fig 2(b)–(d) were assigned according to the subject-specific values reported in the dataset, as summarised in Table 2.

**Fig 2.**
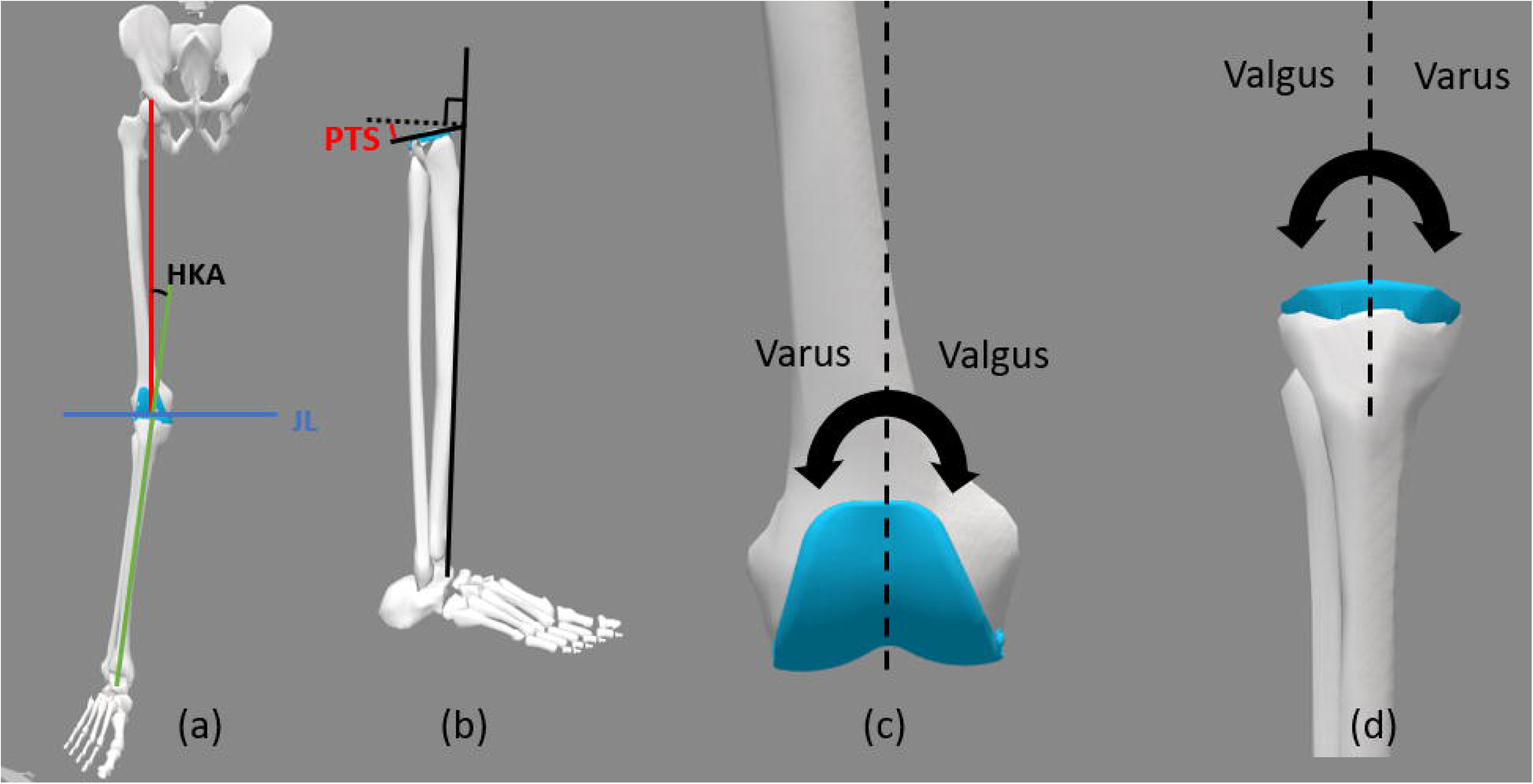
Important angles of TKR components. (a) Hip-Knee-Ankle (HKA) Angle. (b) Posterior Tibial Slope (PTS). (c) Varus-Valgus Alignment of the Femoral Component. (d) Varus-Valgus Alignment of the Tibial Component.

**Table 2.**
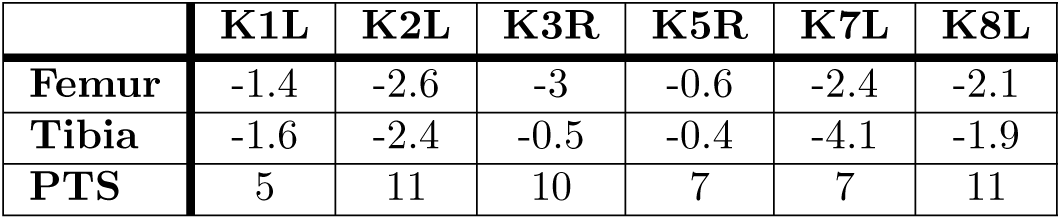
Alignments (in degrees) of replacement component for each subject [38]. Positive values indicate valgus alignment while negative values indicate varus alignment. Positive values for PTS indicate anti-clockwise rotation (see Fig 2).

|  | **K1L** | **K2L** | **K3R** | **K5R** | **K7L** | **K8L** |
|---|---|---|---|---|---|---|
| **Femur** | -1.4 | -2.6 | -3 | -0.6 | -2.4 | -2.1 |
| **Tibia** | -1.6 | -2.4 | -0.5 | -0.4 | -4.1 | -1.9 |
| **PTS** | 5 | 11 | 10 | 7 | 7 | 11 |

### Formulation of the optimal control problem

With the skeletal model described above, an OCP was formulated to simulate the stand-to-squat-to-stand motion. The OCP was solved using OpenSim Moco [28], an open-source tool that uses direct collocation to solve OCPs with musculoskeletal models built in OpenSim. The OCP used in this study is formulated in Eq. (3).

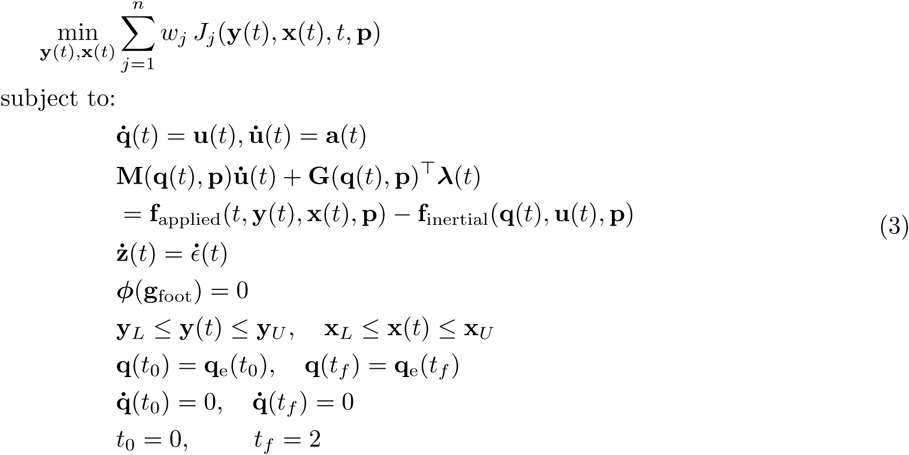

Eq. (3) minimises an integrated cost function over time, where **y** is the state vector, including the generalised coordinates **q** and velocities **u**, and **x** is the control vector associated with the model actuators. The path constraints (**y***_L_*, **y***_U_*, **x***_L_*, and **x***_U_*) enforce realistic joint ranges of motion. The lower-limb coordinate limits were set to 0^◦^–120^◦^ of flexion at both the hip and the knee, and 0^◦^–30^◦^ of dorsiflexion at the ankle [29]. Boundary conditions were imposed so that the joint positions at the start and end of the motion matched the subject-specific upright configuration observed in the experimental data, with zero joint velocities at both endpoints. The auxiliary state **z** tracks ligament strain dynamics through *ɛ̇*, and the kinematic constraint ***ϕ*** is applied to a point on the foot **g**_foot_ to limit foot movement. The weight *w_j_* is the relative contribution of term *J_j_*. With the effort penalty split by actuator, *n* = 11.

The objective function terms employed in this study were defined as follows:

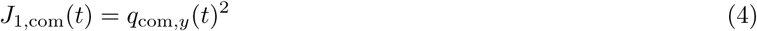

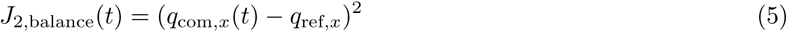

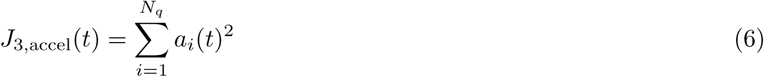

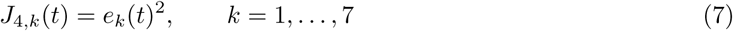

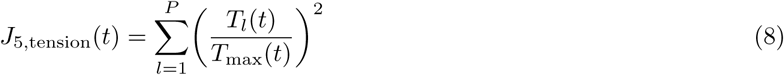

In these expressions, *q*_com*,y*_(*t*) and *q*_com*,x*_(*t*) denote the vertical and anterior-posterior positions of the whole-body centre of mass, and *q*_ref*,x*_ is a fixed anterior-posterior reference position associated with the foot. *a_i_*(*t*) is the acceleration of the *i*^th^ of the *N_q_* generalised coordinates of the model. *e_k_*(*t*) is the excitation of the *k*^th^ torque actuator, bounded between *−*1 and 1. Seven torque actuators were applied to hip flexion/extension, hip adduction/abduction, hip rotation, knee flexion/extension, ankle dorsiflexion/plantarflexion, subtalar inversion/eversion, and lumbar extension. Each has its own weight *w*_4*,k*_. *T_l_*(*t*) is the tension in the *l*^th^ of the *P* ligament components at time *t*, and *T*_max_(*t*) is the maximum ligament tension at that same time point. The outer-level vector **w** therefore has 11 coordinates: *w*_com_, *w*_balance_, *w*_acc_, *w*_tension_, and *w*_4*,k*_ for *k* = 1*, . . . ,* 7. All 11 are constrained to be non-negative and to sum to one.

The centre-of-mass term (*J*_1,com_) is required in this formulation because the boundary conditions constrain the motion to begin and end in an upright posture, and the remaining terms penalise deviation from quiet standing, an objective lacking a term that rewards lowering the centre of mass would be minimised by remaining upright throughout the trial. *J*_1,com_ is therefore required to produce a genuine squatting motion within the optimisation, rather than reflecting a TKR-specific clinical priority.

The balance term (*J*_2,balance_) reflects the postural stability demands of high-flexion tasks, which are important after TKR. Squatting has been identified as one of the most functionally limited and difficult activities reported by TKR patients postoperatively [10].

The acceleration term (*J*_3,accel_) penalises jerky motion and encourages smoother trajectories. It is a standard regulariser in trajectory optimisation [11, 12], but not a TKR-specific term.

Each effort term (*J*_4*,k*_) penalises large excitation of one torque actuator, consistent with reports of persistent quadriceps strength deficits at long-term follow-up after TKR relative to healthy controls [30], which may limit patients’ capacity to sustain high joint torques throughout the squat cycle.

Finally, the ligament tension term (*J*_5,tension_) penalises excessive ligament loading; ligament balance is widely regarded as a critical determinant of TKR outcome and stability [31], so it was included as an objective term in this study. The ligament model itself follows the formulation of Blankevoort et al. [24].

### Bi-level optimisation

Given the cost-function structure in Eqs. 4–8, the corresponding weights must be determined. We therefore employed a bi-level inverse optimal control (IOC) framework. Bi-level IOC has been widely used in human motion research to infer cost-function weights from observed movement [32–35].

In this study, bi-level IOC was used to identify the weight factors associated with post-TKR squat motion. At the inner level, an OCP (Eq. (3)) was solved to generate a squat trajectory for a given set of weight factors **w**. At the outer level, the derivative-free optimisation algorithm PRIMA [33, 36, 37] adjusted these weights to minimise the discrepancy between simulated and observed squat kinematics.

Three IOC settings were considered. In the individualised setting, the cost-function weights were optimised separately for each subject using that subject’s observed squat trajectory. This setting was used to quantify subject-specific reconstruction accuracy and to provide a benchmark for the best achievable agreement under the selected model and objective-function structure. Algorithm 1 outlines this individualised bi-level IOC procedure.

#### Algorithm 1

Individual Bi-Level IOC

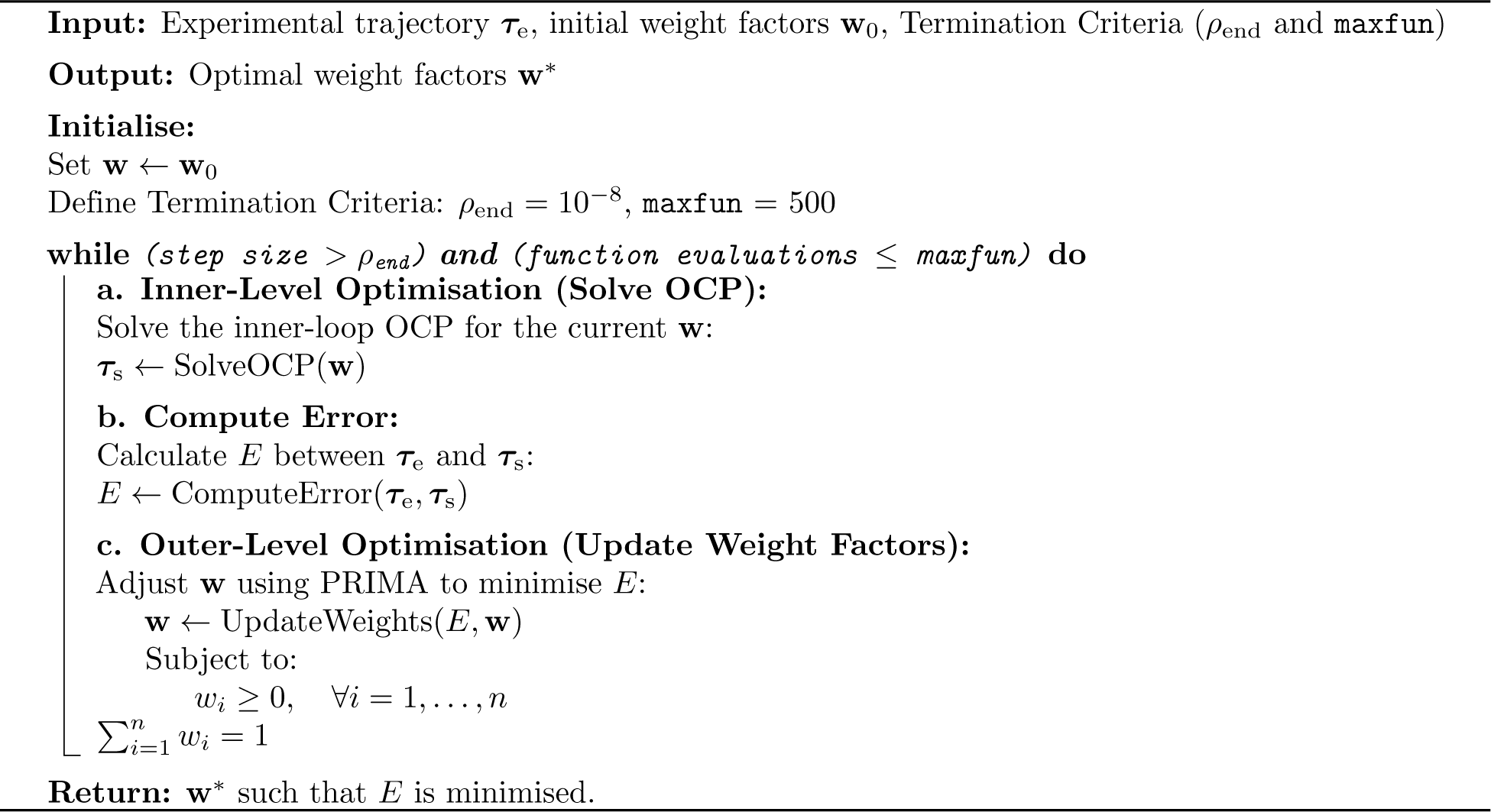

Here, ***τ***_e_ and ***τ***_s_ denote the experimental and simulated lower-limb joint trajectories for hip, knee, and ankle flexion/extension. The discrepancy between the experimental and simulated data was quantified using the apex angle difference and RMSE for each joint:

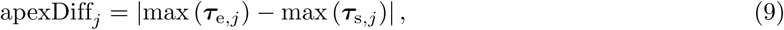

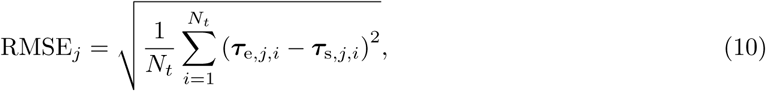

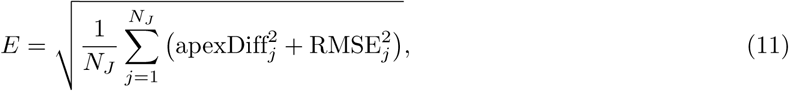

where *j* denotes the joint, *N_t_* is the number of time samples, and *N_J_* = 3 is the number of evaluated joints. The apex angle difference represents the difference in maximum flexion angle between the experimental and simulated trajectories, while RMSE quantifies the average trajectory error across the full squat cycle. This loss treats hip, knee, and ankle flexion equally and does not give extra weight to peak knee flexion, even though that angle is the clinical marker used above.

*ρ*_end_ and maxfun were used as the termination criteria for the upper-level optimisation, representing the tolerance threshold and the maximum number of function evaluations, respectively.

On the other hand, in the group-level setting, a single set of cost-function weights was optimised across all subjects. This setting was used to examine whether one population-level cost function could reproduce the general squat kinematic patterns observed across the available cohort. Because every subject contributes to both the weight identification and the evaluation, this setting is a pooled within-sample fit rather than a predictive test, and is referred to as the in-sample group-level setting throughout this study. Algorithm 2 outlines this group bi-level IOC framework.

#### Algorithm 2

Group Bi-Level IOC

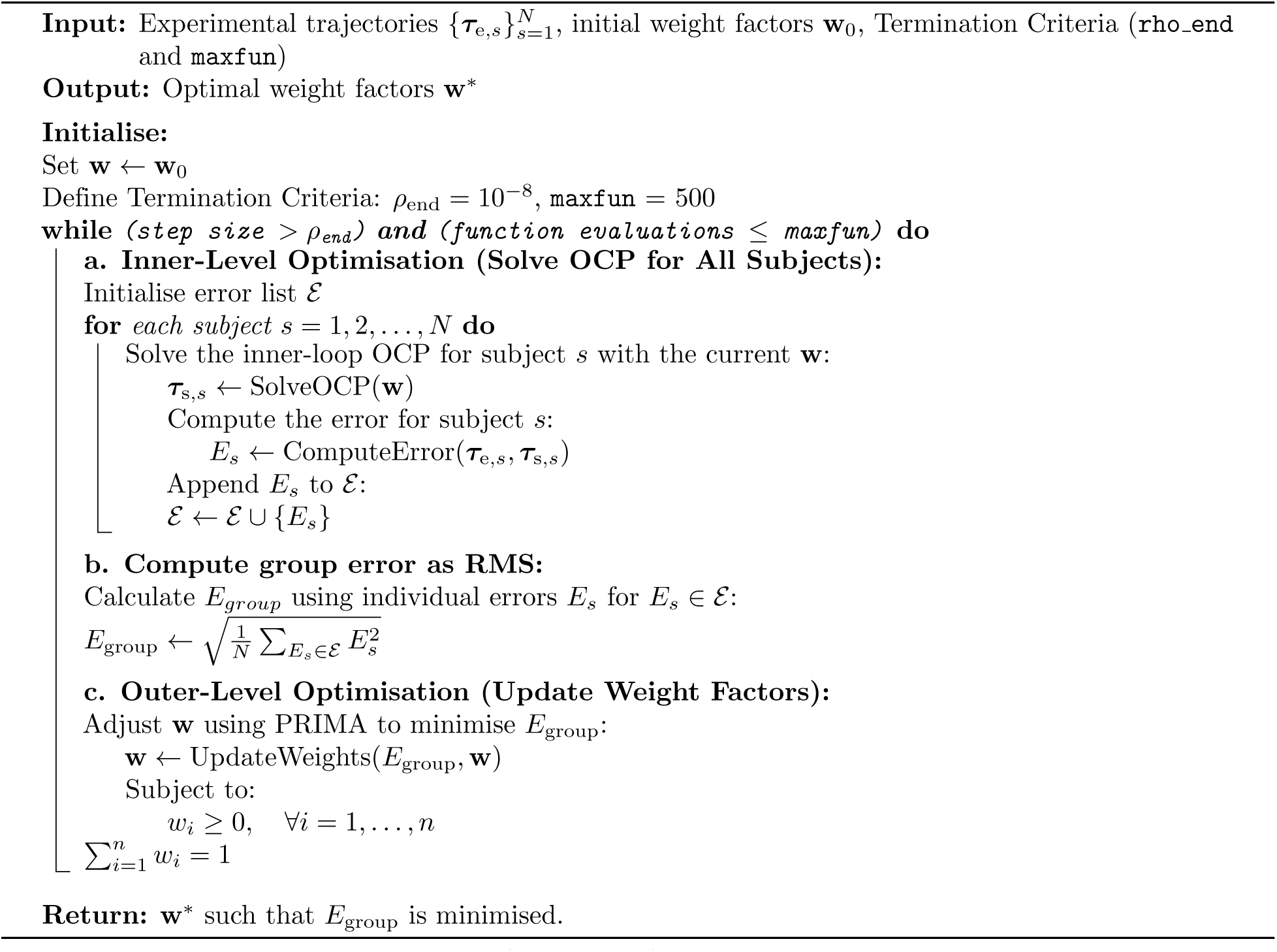

In contrast to the individualised setting (Algorithm 1), the inner-level optimisation was executed separately for each subject using the same weight vector **w**. The subject-specific errors were then aggregated into the set *E* and combined into the group-level error metric, *E*_group_, which was minimised in the outer-level optimisation.

Finally, leave-one-subject-out cross-validation (LOSO-CV) was used to test the shared cost function on subjects who had not entered the fit. For each subject *s*, Algorithm 2 was rerun on the other five, with the same termination criteria, constraints, and starting weights as the in-sample group-level run. That fold produced weights **w***^∗^*. Those weights were then held fixed while the inner OCP (Eq. (3)) was solved for subject *s*, and the simulated trajectory was scored against that subject’s measured squat using Eqs. (9)–(11). Subject *s* therefore did not influence the weights used to simulate their own motion. Outer-level run times were stored for each fold so that the cost of identifying a group-level weight vector could be reported.

### Experimental data

Experimental data were obtained from the CAMS-Knee database [17, 18] and used to estimate cost-function weights and compare observed joint kinematics with simulated trajectories. The dataset included six subjects with TKR (5 male, 1 female; mean age 75 (*±*5) years, weight 89 (*±*13) kg, height 172 (*±*4) cm), with data collected 64–87 months after surgery.

Skin marker trajectories were recorded during squatting. A generic model was scaled and processed using the OpenSim scaling tool and inverse kinematics solver [19] to obtain lower-limb joint angle trajectories. The scaled MSK models were assigned component orientations matching those reported in the dataset [38] to facilitate comparison between simulated and experimental squat kinematics. Replacement component alignments are summarised in Table 2. Negative values indicate varus alignment and positive values indicate valgus alignment, as illustrated in Fig 2.

The code and models used here are openly archived at https://doi.org/10.5281/zenodo.20585083. The archive holds the inner-level Moco problem, the individualised, group-level and cross-validation drivers, the custom objective-term plugin sources, and the OpenSim models. The CAMS-Knee data themselves are not redistributed and remain available from the database under its own data-use agreement.

#### Ethics statement

This study used previously collected human participant data from the CAMS-Knee dataset [17, 18]. No new human participant recruitment, intervention, or data collection was performed for the present secondary analysis. The data were obtained under the CAMS-Knee data-use agreement and were accessed for research purposes on 17 August 2021. The author did not have access to information that could directly identify individual participants during or after accessing the dataset. Ethical approval and informed consent procedures for the original CAMS-Knee data collection are reported in the dataset documentation and associated publications [17, 18].

Access was granted after a formal data request, which the CAMS-Knee data providers reviewed and approved. The present work is a secondary analysis of previously collected, de-identified data, with no new recruitment, intervention, or data collection. No additional institutional ethical review was sought.

## Results

The bi-level inverse optimal control framework successfully recovered cost-function weights that reproduced post-total knee replacement (TKR) squat kinematics across all six subjects from the CAMS-Knee dataset. Agreement between simulated and experimental trajectories was quantified using four complementary metrics: apex joint angle difference, defined as the difference in peak flexion between simulated and experimental trajectories; root-mean-square error (RMSE); normalised RMSE (nRMSE), scaled to each joint’s experimental range of motion; and Pearson correlation coefficient (*r*) for the hip, knee, and ankle flexion/extension trajectories. The correlation coefficient tracks the shape and phasing of a trajectory, not its amplitude. A high value can sit alongside a large apex or RMSE error, as it does at the knee in the in-sample group-level setting (Table 4). All four metrics are therefore reported together. Results are presented separately for the individualised subject-specific setting, the in-sample group-level setting, and the leave-one-subject-out cross-validation. All simulations satisfied the non-negativity and normalisation constraints on the weight factors and terminated within the prescribed limits of the upper-level optimisation.

### Individual cost function results

In the individualised setting, the optimised weights produced squat trajectories that closely reproduced the experimental kinematics across all six subjects. The simulated and experimental trajectories for the three lower-body degrees of freedom in the sagittal plane are shown in Fig 3.

**Fig 3.**
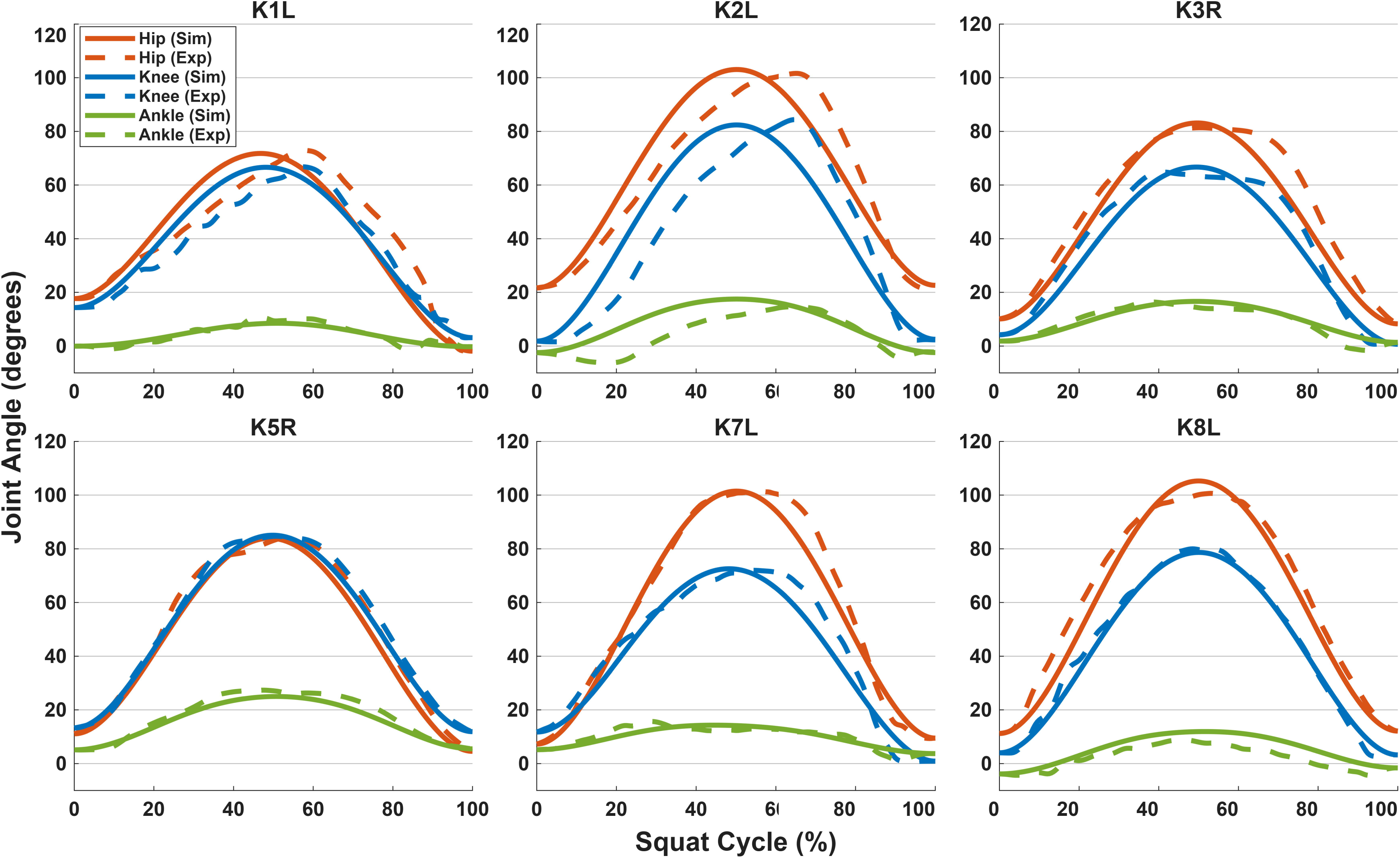
Trajectories for the individual weight setting. Solid lines represent simulated trajectories and dashed lines represent experimental trajectories for the hip, knee, and ankle flexion/extension angles.

Subject-specific optimisation of the cost-function weights enabled the model to closely match the experimental squat kinematics for each participant. Apex joint angle differences between simulated and experimental trajectories ranged from 0.02^◦^ at the hip of subject K7L to 4.65^◦^ at the hip of subject K8L. Averaged across all subjects, these differences were 1.55 (*±*1.66)^◦^ for the hip, 1.05 (*±*0.78)^◦^ for the knee, and 1.98 (*±*1.09)^◦^ for the ankle, giving an overall mean of 1.53 (*±*1.18)^◦^. RMSE across the full squat cycle averaged 6.87 (*±*2.41)^◦^ at the hip, 5.71 (*±*3.69)^◦^ at the knee, and 2.87 (*±*1.85)^◦^ at the ankle. When scaled to each joint’s range of motion, nRMSE was 8.56 (*±*3.44)% for the hip, 7.84 (*±*4.40)% for the knee, and 17.04 (*±*9.26)% for the ankle. Pearson correlation coefficients exceeded 0.9 for every joint and subject, with mean values of 0.97 (*±*0.03) for both the hip and knee and 0.93 (*±*0.06) for the ankle. Summary metrics for the individualised setting are presented in Table 3.

**Table 3.**
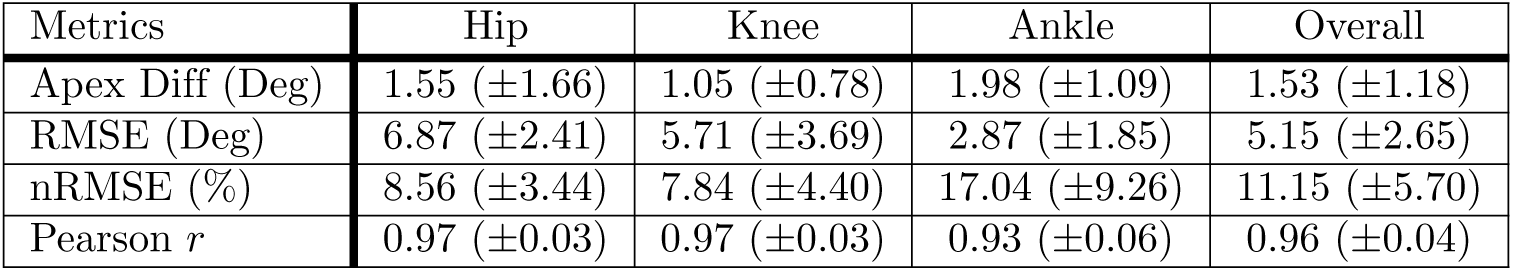
Metrics averaged across all subjects for the individualised setting. Results are reported for each joint (hip, knee, and ankle) as well as overall averages. Values are presented as mean (*±*standard deviation).

### In-sample group cost function results

In the in-sample group-level setting, a single group-level weight set was optimised across all subjects simultaneously. The simulated and experimental trajectories for this setting are shown in Fig 4.

**Fig 4.**
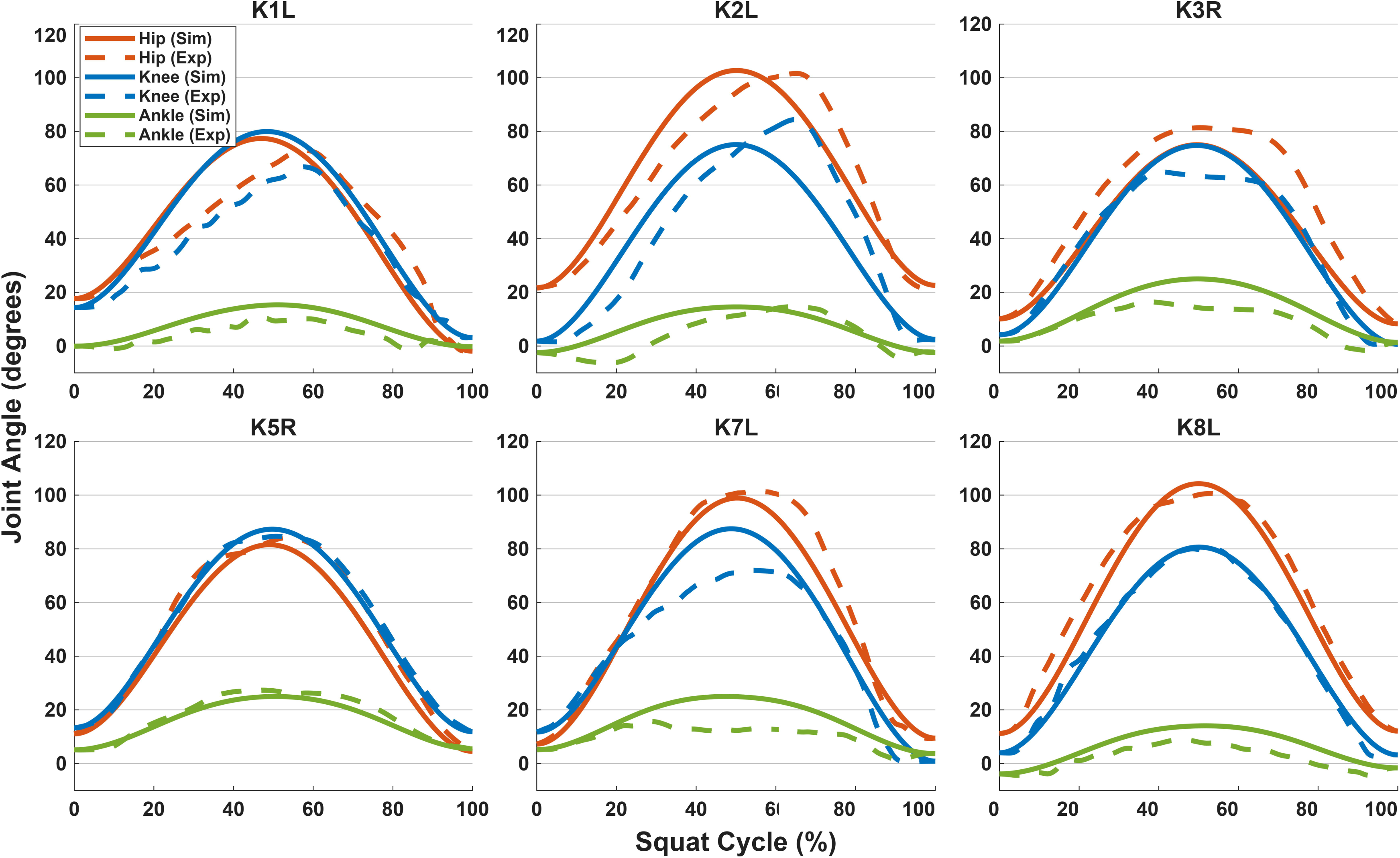
Trajectories for the in-sample group weight setting. Solid lines represent simulated trajectories and dashed lines represent experimental trajectories for the hip, knee, and ankle flexion/extension angles.

The in-sample group-level weights produced trajectories that followed the general pattern of the experimental squat motion within the available cohort, although with larger deviations than the individualised setting. Apex joint angle differences ranged from 0.14^◦^ at the ankle of subject K2L to 15.45^◦^ at the knee of subject K7L. Mean differences across subjects were 3.46 (*±*1.81)^◦^ for the hip, 8.52 (*±*5.82)^◦^ for the knee, and 5.12 (*±*3.53)^◦^ for the ankle, resulting in an overall mean of 5.70 (*±*3.72)^◦^. RMSE averaged 8.03 (*±*2.58)^◦^ for the hip, 7.17 (*±*4.75)^◦^ for the knee, and 5.05 (*±*1.77)^◦^ for the ankle. Normalised RMSE was 10.07 (*±*3.98)% for the hip, 10.08 (*±*6.76)% for the knee, and 32.44 (*±*14.00)% for the ankle. Pearson correlation coefficients remained high, averaging 0.97 (*±*0.03) for both the hip and knee and 0.92 (*±*0.07) for the ankle. Summary metrics for the in-sample group-level setting are reported in Table 4.

**Table 4.** Metrics averaged across all subjects for the in-sample group-level setting. Results are reported for each joint (hip, knee, and ankle) as well as overall averages. Values are presented as mean (*±*standard deviation).

| Metrics | Hip | Knee | Ankle | Overall |
|---|---|---|---|---|
| Apex Diff (Deg) | 3.46 (±1.81) | 8.52 (±5.82) | 5.12 (±3.53) | 5.70 (±3.72) |
| RMSE (Deg) | 8.03 (±2.58) | 7.17 (±4.75) | 5.05 (±1.77) | 6.75 (±3.03) |
| nRMSE (%) | 10.07 (±3.98) | 10.08 (±6.76) | 32.44 (±14.00) | 17.53 (±8.25) |
| Pearson *r* | 0.97 (±0.03) | 0.97 (±0.03) | 0.92 (±0.07) | 0.95 (±0.04) |

### Cross-validated group cost function results

In the leave-one-subject-out cross-validation, each subject was simulated using group-level weights identified from the remaining five subjects only. The simulated and experimental trajectories for this setting are shown in Fig 5.

**Fig 5.**
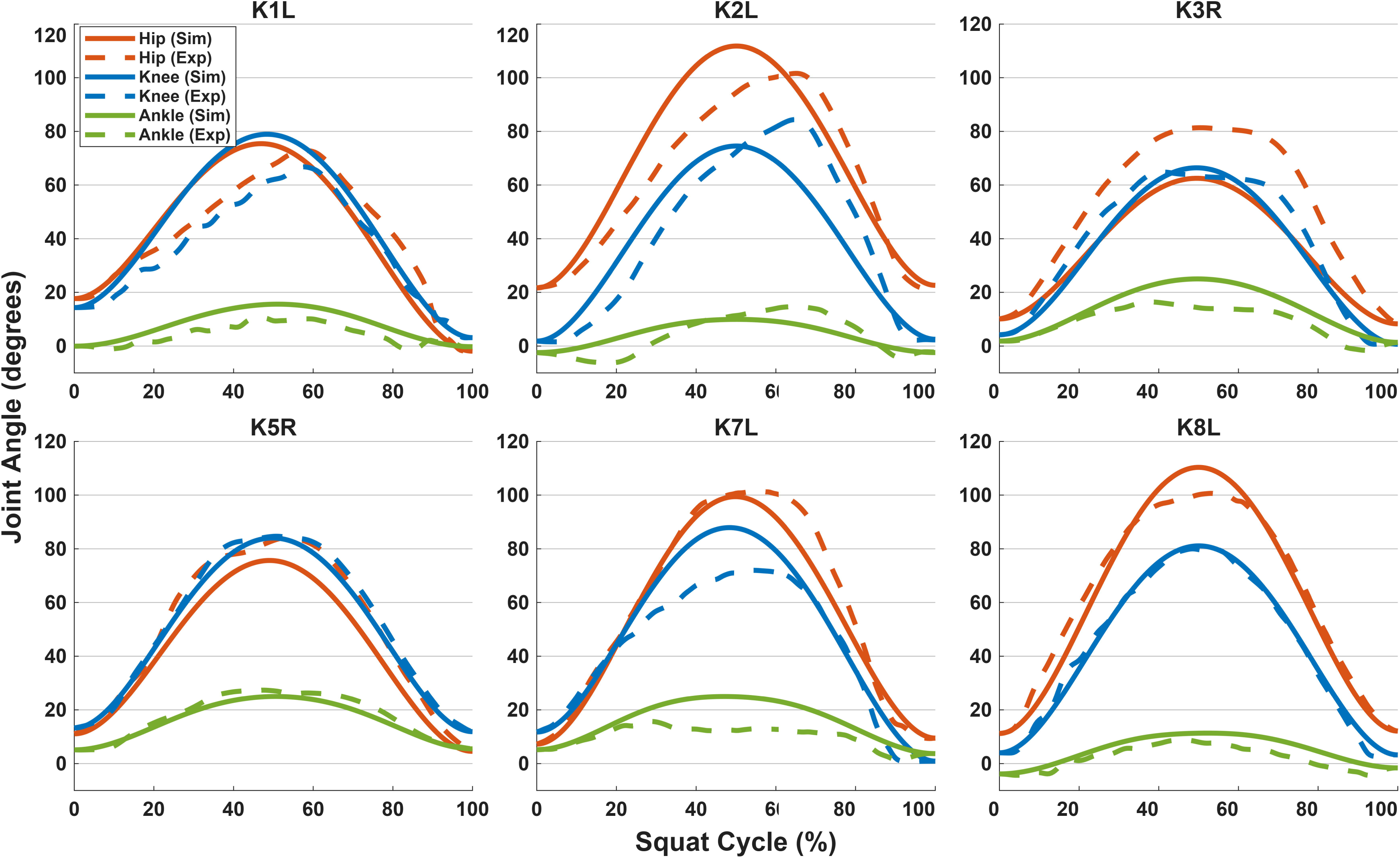
Trajectories for the cross-validated group weight setting. Solid lines represent simulated trajectories and dashed lines represent experimental trajectories for the hip, knee, and ankle flexion/extension angles. Each subject was simulated using group-level weights identified from the remaining five subjects.

Cross-validated apex joint angle differences ranged from 0.64^◦^ at the knee of subject K5R to 18.87^◦^ at the hip of subject K3R. Mean differences across subjects were 8.66 (*±*6.15)^◦^ for the hip, 6.86 (*±*6.62)^◦^ for the knee, and 5.48 (*±*2.92)^◦^ for the ankle, giving an overall mean of 7.00 (*±*5.23)^◦^. RMSE averaged 10.05 (*±*4.51)^◦^ for the hip, 7.31 (*±*4.46)^◦^ for the knee, and 4.71 (*±*1.86)^◦^ for the ankle. Normalised RMSE was 12.74 (*±*6.69)% for the hip, 10.26 (*±*6.27)% for the knee, and 30.25 (*±*14.39)% for the ankle. Pearson correlation coefficients averaged 0.97 (*±*0.03) for both the hip and knee and 0.92 (*±*0.07) for the ankle. Summary metrics for the cross-validated setting are reported in Table 5.

**Table 5.** Metrics averaged across all subjects for the leave-one-subject-out cross-validation. Results are reported for each joint (hip, knee, and ankle) as well as overall averages. Values are presented as mean (*±*standard deviation).

| Metrics | Hip | Knee | Ankle | Overall |
|---|---|---|---|---|
| Apex Diff (Deg) | 8.66 (±6.15) | 6.86 (±6.62) | 5.48 (±2.92) | 7.00 (±5.23) |
| RMSE (Deg) | 10.05 (±4.51) | 7.31 (±4.46) | 4.71 (±1.86) | 7.36 (±3.61) |
| nRMSE (%) | 12.74 (±6.69) | 10.26 (±6.27) | 30.25 (±14.39) | 17.75 (±9.12) |
| Pearson *r* | 0.97 (±0.03) | 0.97 (±0.03) | 0.92 (±0.07) | 0.95 (±0.04) |

Comparison with the in-sample group-level results in Table 4 shows that the accuracy lost by evaluating the cost function on the held-out subjects was small on average but not evenly distributed among joints. Overall normalised RMSE increased from 17.53% to 17.75%, overall RMSE from 6.75^◦^ to 7.36^◦^, and the overall Pearson correlation coefficient was unchanged at 0.95. At the knee and ankle the two settings were closely comparable, with differences in both directions. Normalised RMSE was 0.18 percentage points higher under cross-validation at the knee but 2.19 percentage points lower at the ankle, and the mean apex joint angle difference was 1.66^◦^ lower under cross-validation at the knee but 0.36^◦^ higher at the ankle. The difference was concentrated at the hip, where the mean apex joint angle difference increased from 3.46^◦^ to 8.66^◦^ and normalised RMSE increased from 10.07% to 12.74%.

Once a group-level weight vector has been identified, simulating a subject who did not contribute to that identification requires a single inner-level OCP solution. Across the six held-out subjects this took 671 (*±*261) s, ranging from 371 s to 999 s. Identifying the weights is considerably more expensive but is performed once. Each cross-validation fold required between 61 and 138 outer-level function evaluations (mean 77.8), and each evaluation required one inner-level OCP solution for each of the five training subjects, all solved sequentially. The cumulative inner-level solution time was 70.8 h per fold on average, ranging from 49.1 h to 111.7 h. Although weight identification is more than a hundred times slower than application, once the weights exist, using them on a new subject will cost one inner-level OCP.

### Comparison between cost function results

Fig 6 provides a direct comparison of the four metrics between the individualised, in-sample group-level, and cross-validated group-level cost functions.

**Fig 6.**
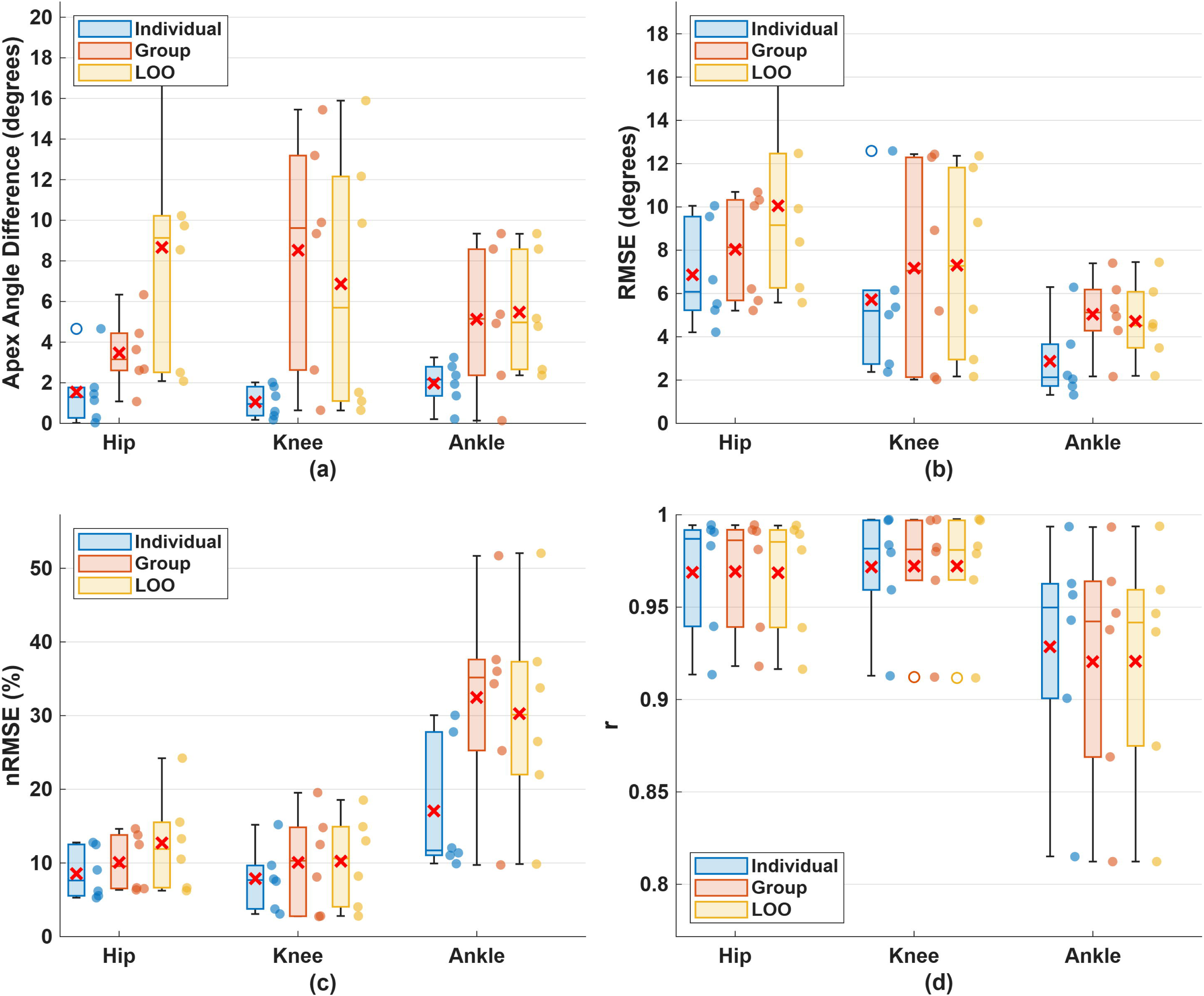
Box plots comparing result metrics between the individualised, in-sample group-level, and cross-validated cost functions. Scatter points adjacent to each box indicate the raw metric values for each subject, while red x marks represent the mean values. Subplots display (a) apex joint angle difference, (b) RMSE, (c) normalised RMSE, and (d) Pearson’s correlation coefficient, with results presented for the hip, knee, and ankle joints.

The individualised cost function yielded lower mean apex joint angle differences than the in-sample group-level cost function across all three joints. The difference was particularly evident at the knee, where the mean apex difference decreased by 87.7% in the individualised setting. Reductions were also observed at the hip and ankle. A comparable pattern was found for RMSE, which was lower in the individualised setting by 14.4% at the hip, 20.4% at the knee, and 43.2% at the ankle. Normalised RMSE followed the same trend, showing reductions of 15.0% at the hip, 22.2% at the knee, and 47.5% at the ankle. In contrast, Pearson correlation coefficients remained high and largely comparable between the two settings for the hip and knee joints, with only a small difference observed at the ankle (0.93 in the individualised setting versus 0.92 in the in-sample group-level setting).

The same pattern held against the cross-validated setting. Relative to the cross-validated cost function, the individualised setting produced apex joint angle differences lower by 82.1% at the hip, 84.7% at the knee, and 63.9% at the ankle, and normalised RMSE lower by 32.8%, 23.6%, and 43.7% respectively.

Paired effect sizes for all three pairwise comparisons are reported in Table 6, computed across the six subjects using each subject’s mean value over the three joints. Paired Cohen’s *d_z_* was the mean of the six subject-level differences divided by their standard deviation. 95% confidence intervals used the non-central *t* distribution. No hypothesis test is reported. The sample size is set by the cohort rather than chosen, so neither a significance test nor a power calculation would be informative, and we are not aware of a minimum difference of interest for these metrics in post-TKR squat simulation. The individualised setting differed from both group-level settings by a large margin, with Cohen’s *d_z_* between *−*0.87 and *−*1.90 across the three error metrics, and the confidence intervals for the apex joint angle difference excluded zero in both comparisons. The two group-level settings differed only marginally on every metric, and were closest on normalised RMSE, where *d_z_* was *−*0.09 with a 95% confidence interval from *−*0.89 to 0.71. With six subjects these intervals are wide. Failing to find a difference between the in-sample and leave-one-out settings is not the same as showing they are equal; a moderate effect would still be compatible with these data.

**Table 6.** Paired effect sizes between the three settings. Cohen’s *d_z_* with 95% confidence intervals, computed across the six subjects using each subject’s mean value over the three joints. Negative values indicate that the first-named setting produced the lower value of the metric.

| Metric | Individual vs in-sample | Individual vs LOSO-CV | In-sample vs LOSO-CV |
|---|---|---|---|
| Apex Diff | −1.18 [−2.22, −0.08] | −1.90 [−3.27, −0.48] | −0.67 [−1.54, 0.25] |
| RMSE | −0.91 [−1.84, 0.09] | −1.08 [−2.07, −0.02] | −0.57 [−1.41, 0.33] |
| nRMSE | −0.93 [−1.88, 0.08] | −0.87 [−1.80, 0.11] | −0.09 [−0.89, 0.71] |
| Pearson *r* | 0.58 [−0.32, 1.43] | 0.75 [−0.20, 1.64] | 0.24 [−0.58, 1.04] |

### Recovered weight factors

The recovered weight factors for each subject in the individualised setting and the single set of weights for the in-sample group-level setting are shown in Fig 7. The centre-of-mass height term (*w*_com_) and the balance term (*w*_balance_) were consistently among the dominant terms, together accounting for more than half of the total weight in most cases. The balance term showed the greatest inter-subject variability, ranging from 0.0950 to 0.4158. The ligament tension term (*w*_tension_) and acceleration term (*w*_acc_) remained small, on the order of 10*^−^*^3^ and 10*^−^*^4^. Those two coordinates share the unit simplex with the centre-of-mass, balance, and seven per-actuator effort weights, so a small value is not by itself evidence that the term is irrelevant. The effort-related terms exhibited moderate subject-specific variation across coordinates.

**Fig 7.**
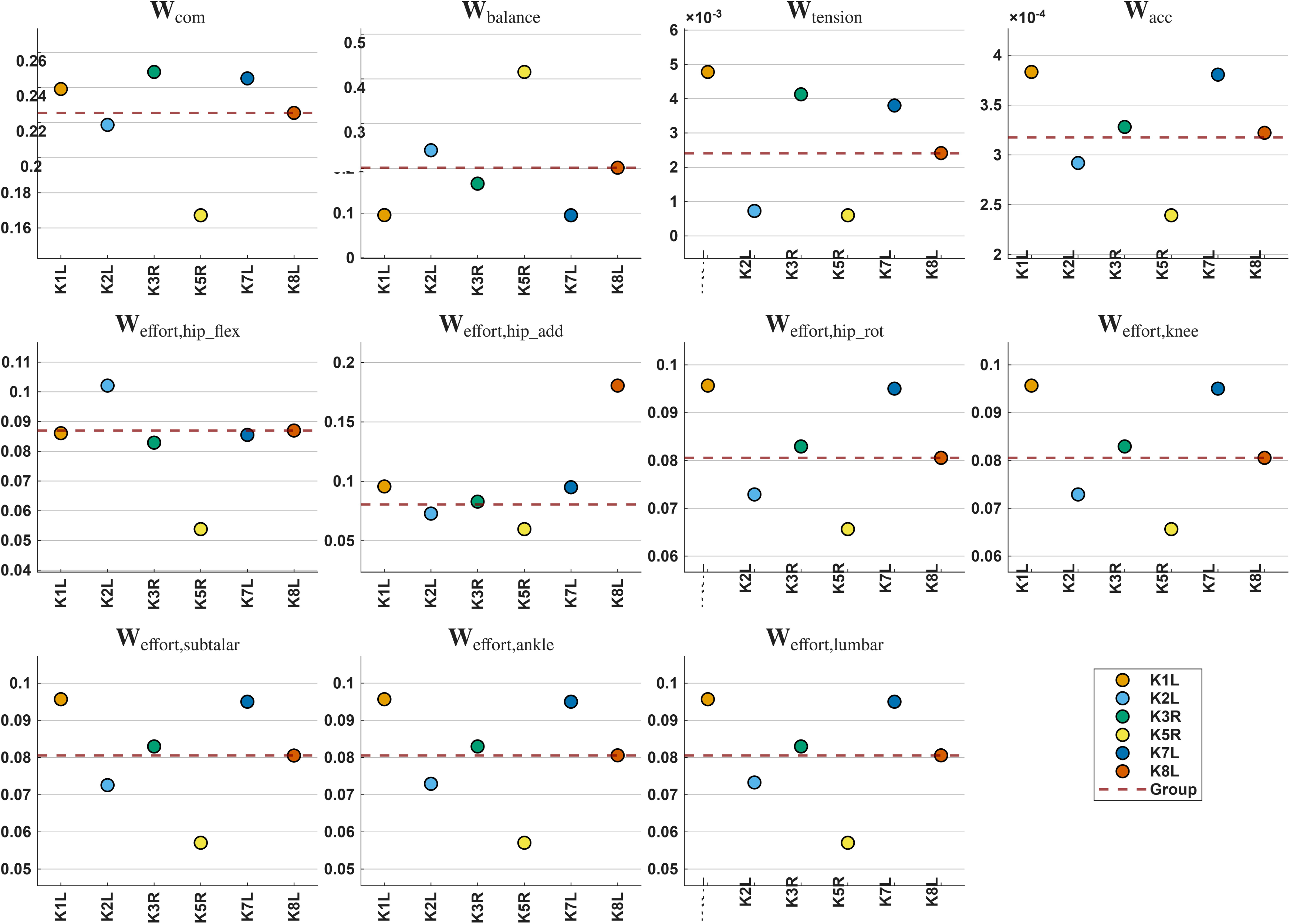
Recovered weight factors for each subject in the individualised setting. Each subplot shows the weight factor for one objective term used in the OCP (Eqs 4 to 8), with one point per subject. The dashed horizontal line marks the corresponding in-sample group-level weight. Vertical scales differ between subplots. Subscripts on the effort weights indicate the coordinate to which the torque actuator is applied; for example, w_effort,hip flex_ is the weight factor for the term that minimises the activation of the torque actuator in the hip flexion/extension coordinate.

The group-level weights identified in each cross-validation fold are shown in Fig 8, together with the in-sample group-level value of each term. Every fold started from the same weights as the in-sample group-level run, so the plot shows how far the solution moved after one subject was dropped, not six independent searches. The movement was small for the dominant terms: the centre-of-mass and balance weights differed from their in-sample group-level values by no more than 1.1% in any fold, and no weight differed by more than 0.0022 in absolute value. The largest proportional change occurred in the ligament tension term, which is small in absolute value and differed from its in-sample group-level value by between 1.0% and 12.2% depending on the fold. The recovered weights were therefore only weakly affected by the exclusion of any single subject, whereas the resulting simulated kinematics were not. For subject K3R, every weight lay within 1.0% of its in-sample group-level value, yet the hip apex joint angle difference increased from 6.34^◦^ to 18.87^◦^, and across folds the change in hip kinematics was not ordered by the size of the change in weights.

**Fig 8.**
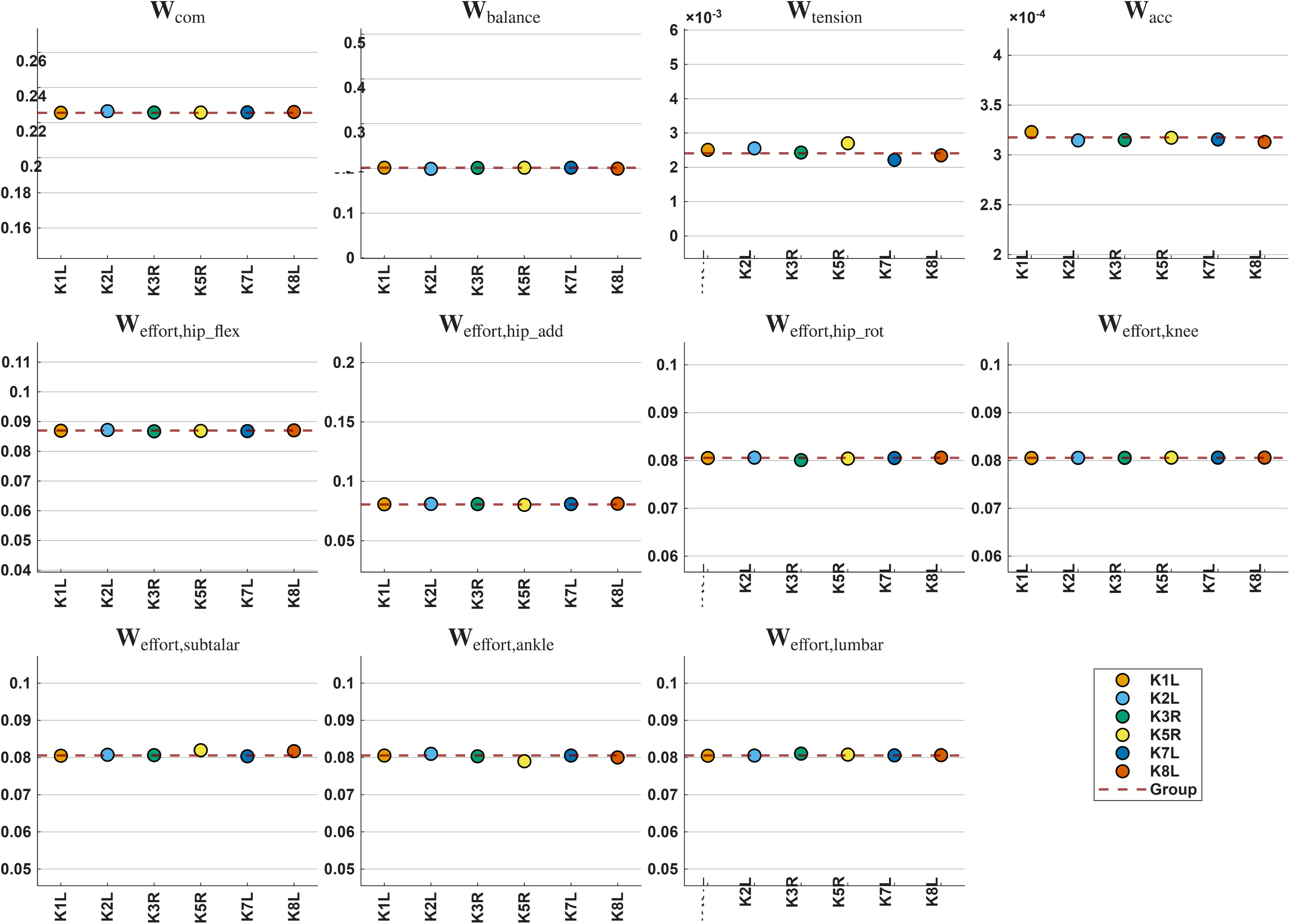
Group-level weight factors recovered in each cross-validation fold. Each subplot represents the weight factor for one objective term, with one point per fold, labelled by the subject held out in that fold. The dashed horizontal line marks the corresponding in-sample group-level weight. Note that the vertical scales differ between subplots and are expanded to resolve differences that are small in absolute terms.

## Discussion

This study developed and preliminarily evaluated a bi-level inverse optimal control framework for simulating post-TKR squat kinematics in three settings. In the individualised setting, subject-specific optimisation produced low apex joint angle differences, RMSE, and nRMSE values, establishing an upper bound on the accuracy achievable under the present model and objective-function structure. The in-sample group-level setting, in which a single set of weights was optimised across all subjects, yielded moderately higher errors. On the pooled metrics the extra error was small. Under leave-one-subject-out cross-validation, overall normalised RMSE rose from 17.53% to 17.75%, overall RMSE from 6.75^◦^ to 7.36^◦^, and the overall Pearson correlation coefficient was unchanged at 0.95. The pooled figures do not show where that change sits. These findings extend our preliminary simulation work [16] by incorporating both femoral and tibial component alignments, a 3-DOF knee joint with ligament dynamics, and comparison with experimental squat data from TKR patients.

Leave-one-subject-out tests the weights, not a preoperative patient. In each fold, only the cost-function weights were withheld. The held-out subject’s scaled model, implant alignment values, and the boundary conditions of the optimal control problem were all derived from their own postoperative data, none of which would be available before surgery, and that subject was drawn from the same six-patient cohort rather than from an independent population. The cross-validation establishes that the identified cost function is not specific to the subjects used to derive it. Using the method before surgery would still require a means of specifying the subject-specific model and its boundary conditions from preoperative information alone.

Averaging across joints (Overall columns in Tables 4 and 5) hides where the extra error sits. At the knee and ankle the in-sample and cross-validated settings were closely comparable, with differences in both directions, and at the knee the cross-validated setting produced the smaller mean apex joint angle difference. The loss was concentrated at the hip, where the mean apex joint angle difference rose from 3.46^◦^ to 8.66^◦^ and normalised RMSE from 10.07% to 12.74%. That mean mixes different subjects. Under leave-one-out the held-out hip was under-flexed in K3R and over-flexed in K2L. That pattern is consistent with an identification loss that does not privilege the knee. None of the confidence intervals comparing the two group-level settings excluded zero, so this difference is moderate in magnitude rather than statistically resolved at the present sample size.

The simulation results showed a level of agreement with experimental kinematics that was broadly comparable to related studies, although direct comparisons should be interpreted cautiously because of differences in study populations, movement tasks, and modelling approaches. For example, Lee *et al.* [13] used a Gaussian process regression model to predict joint angles during smith machine squats in healthy young subjects, reporting nRMSE values between 3.8% and 17% and Pearson correlation coefficients (*r*) ranging from 0.984 to 0.999 for the knee joint. In the present study, the individualised setting produced knee nRMSE values ranging from 3.07% to 15.19% and *r* values from 0.91 to 0.99. Under cross-validation the corresponding knee nRMSE values ranged from 2.80% to 18.55%. These values fall within a comparable range, particularly considering that the present study involved free squatting in older TKR patients rather than constrained smith machine squats in healthy participants.

Other studies have reported lower errors for knee flexion-extension prediction in gait and running tasks. Renani *et al.* [39] and Wouda *et al.* [14] reported RMSE values of 3.3^◦^ and below 5^◦^, respectively, in healthy participants. Although these errors are lower than the mean knee RMSE of 5.71^◦^ observed in the individualised setting of the present study, squat motion involves a larger knee range of motion and greater functional demand than level gait or treadmill running. In addition, the TKR population is expected to show greater inter-subject variability because of differences in implant alignment, soft-tissue properties, strength, pain avoidance, and movement strategy.

Tan *et al.* [15] investigated knee joint kinematics during sit-to-stand and stand-to-sit motions, which are mechanically related to the descent and ascent phases of squatting. They reported RMSE values between 7.27^◦^ and 9.30^◦^, nRMSE values from 8.68% to 10.86%, and Pearson correlation coefficients greater than 0.99. In comparison, the present study obtained mean knee RMSE and nRMSE values of 5.71^◦^ and 7.84%, respectively, in the individualised setting, 7.17^◦^ and 10.08%, respectively, in the in-sample group-level setting, and 7.31^◦^ and 10.26%, respectively, under cross-validation. These comparisons suggest that the proposed simulation framework can reproduce post-TKR squat kinematics with errors that are within the range of related movement-prediction studies, while using a physics-based optimisation framework rather than a purely data-driven model.

Although apex joint angle differences are less frequently reported, they are particularly relevant in the present context because peak flexion is closely related to functional squat depth after TKR. Tan *et al.* [15] reported apex knee angle errors ranging from 5.09^◦^ to 6.89^◦^ during sit-to-stand and stand-to-sit tasks. The average apex knee joint angle difference in the present study was 1.05^◦^ in the individualised setting, 8.52^◦^ in the in-sample group-level setting, and 6.86^◦^ under cross-validation, the last of these falling within the range reported by Tan *et al.* The individualised result therefore compares favourably with previous reports, whereas the in-sample group-level result highlights the additional difficulty of reproducing subject-specific squat depth using a single population-level cost function.

The paired effect sizes split that comparison into two steps. Switching from subject-specific weights to shared weights is a large effect (Cohen’s *d_z_* between *−*0.87 and *−*1.90 on the three error metrics), and the confidence intervals for apex angle difference exclude zero in both group-level comparisons. Switching from an in-sample shared cost function to one applied to a left-out subject is a small effect on every metric, and smallest on nRMSE. Sharing the cost function across patients therefore costs more accuracy than then using that shared function on someone who was not in the fit. The second step is the one that matters if postoperative motion will not be available.

The recovered weight factors suggest that a single universal cost function may be insufficient to capture the full variability of post-TKR squat strategies. The centre-of-mass height term (*w*_com_) and balance term (*w*_balance_) were consistently dominant, indicating that lowering the body while maintaining balance was central to reproducing the motion, but the balance term varied considerably between subjects, from 0.0950 to 0.4158, and the effort-related terms also showed subject-specific variation across coordinates. The cross-validation folds limit how far those values can be read as a description of strategy. Excluding any one subject changed the recovered weights very little yet changed the simulated kinematics of that subject appreciably, and not in proportion. For subject K3R every weight lay within 1.0% of its in-sample group-level value while the hip apex joint angle difference increased from 6.34^◦^ to 18.87^◦^, whereas the fold with the largest weight change produced a smaller kinematic change. Weights that differ only slightly can therefore correspond to appreciably different predicted movements, at least at the hip, consistent with the known difficulty of uniquely identifying weights in weighted-sum inverse optimal control [40].

The ankle nRMSE is the largest of the three joints in both group-level settings, at 32.44% in sample and 30.25% under cross-validation. This is largely a normalisation effect. Mean experimental ankle excursion during the squat was 16.8^◦^, against 72.0^◦^ at the knee and 81.9^◦^ at the hip, so a given absolute error is scaled by a denominator roughly four times smaller. In absolute terms the ankle carried the smallest error of the three joints in every setting, averaging 5.05^◦^ in sample and 4.71^◦^ under cross-validation. Ankle dorsiflexion range constrains squat depth and strategy [41], and the plantarflexors controlling it act across both the ankle and, for gastrocnemius, the knee [42]. Because the model is actuated by joint torques rather than muscles, it cannot represent these contributions, nor distinguish a genuine ankle strategy from a torque distribution that reproduces the same kinematics. Three things are specific to this joint. The ankle was not operated on, but it still takes up the altered mechanics of the replaced knee. Its foot and ankle segments come from the generic model, scaled to each subject, so the geometry is not subject-specific. Unlike the knee, the ankle has no passive soft-tissue elements, so passive resistance to dorsiflexion is not modelled; it enters only as a bound on the coordinate. Compensatory strategies at joints adjacent to the replaced knee have been reported after TKR [43], and evaluating whether the framework reproduces them would require muscle-level actuation.

Simulated trajectories in all three settings reproduced squat depth and overall shape more closely than the timing of the descent-to-ascent transition. Several experimental trials were asymmetric, but the simulations were close to symmetric, including in the individualised setting. That is more likely a property of the objective than of the particular weights. The cost function is a weighted sum with a single weight vector applied over the whole trial, so the relative priority given to lowering the centre of mass, maintaining balance and limiting actuator effort is forced to be the same during descent and ascent. Descent and ascent are not mechanically equivalent [44], and a static weight vector cannot express a control priority that changes between them. Phase-dependent or time-varying weights [45] would be the natural way to test this explanation, at the cost of a larger identification problem and a correspondingly greater risk of overfitting at this sample size.

The model uses joint torque actuators rather than muscles. That keeps the inner OCP tractable, but omits co-contraction, activation dynamics, and metabolic cost [46], so compensatory recruitment after TKR cannot be verified [47, 48]. Ligament stiffness, reference strain, and maximum actuator torques were taken from the literature and held the same for every subject. Soft-tissue tension after TKR varies with surgical balance [49]. Modelled post-TKR kinematics also change when collateral stiffness is perturbed, although the shift from residual strain-estimation error is small relative to implant-position error [50]. *J*_5,tension_ uses those shared parameters.

The leave-one-out folds only change the weight vector. Model parameters, boundary conditions, and the set of objective terms were not varied. Weight vectors differing by at most 0.0022 still changed one subject’s hip apex error by more than 12^◦^. PRIMA is a local search method, and all optimisations reported here were initialised from the same weight vector, so the recovered weights may represent local rather than global optima.

Despite these limitations, the framework provides a proof of concept for dynamic functional simulation of post-TKR squat. The individualised setting is an upper-bound benchmark, and the cross-validated setting is an out-of-sample estimate within this cohort. These conclusions rest on six patients from one dataset, one implant setting, and one task, so the confidence intervals are wide.

Future work should first validate the method in an independent cohort, ideally spanning more than one implant design and surgical protocol, and in a larger sample. A designed sensitivity analysis using multi-start identification and systematic parameter perturbation is required to establish how robust the recovered weights and the resulting kinematics are, and would also address the weak identifiability of hip kinematics observed here. Phase-dependent or time-varying objective terms may improve reproduction of the descent-to-ascent timing asymmetry, and global search strategies could reduce the risk of convergence to local minima. Extending the model to muscle-driven actuators [51] would improve biomechanical realism and allow physiologically motivated objectives, such as muscle effort or metabolic energy expenditure, to be investigated. Finally, applying the approach to larger TKR cohorts and exploring clustering based on recovered weight distributions or movement strategies may support more robust population-level cost functions for future preoperative functional simulation.

## Conclusion

This study demonstrates the feasibility of combining musculoskeletal simulation with bi-level inverse optimal control to reproduce post-total knee replacement squat kinematics. The individualised cost-function setting provided accurate subject-specific reconstruction and served as a benchmark for the best achievable agreement under the proposed model and objective-function structure. The in-sample group-level setting produced higher errors but preserved the overall squat motion pattern across subjects. Leave-one-subject-out cross-validation showed that this shared cost function transferred to subjects that had not contributed to its identification with little further loss of accuracy overall, giving an overall normalised RMSE of 17.75% against 17.53% in sample and an unchanged Pearson correlation coefficient of 0.95, although that loss was concentrated at the hip rather than distributed across joints.

The framework generated the full stand-to-squat-to-stand cycle in a single optimisation process without prescribing a target squat depth. The resulting depth follows from the model dynamics, the boundary conditions, and the relative weighting of the cost-function terms, including a term that explicitly rewards lowering of the centre of mass. Agreement inside one six-patient cohort is not the same as predicting squat kinematics before surgery. Larger and independent data are needed before the method could be used for planning.

## Acknowledgments

The author thanks the CAMS-Knee investigators and data providers for providing access to the dataset used in this study.

